# Differentiation potential of fish fins –Effective utilization of the fins as food wastes–

**DOI:** 10.1101/2020.02.29.950386

**Authors:** Yusuke Tsuruwaka, Eriko Shimada

**Author notes:** Corresponding author: Yusuke Tsuruwaka, E-mail address, E-mail address, Eriko Shimada.

## Abstract

Fish cells are largely affected by their culture media. Fibroblast-like cells obtained from the fish fins can differentiate to various kinds of cells such as skeletal muscle-like, neurofilaments and adipocytes. Our results suggest that the fins which are usually discarded as food wastes may practically applied to the clean meat technology.

## Introduction

*In vitro* meat production technology, also known as the cell culturing technology of ‘clean meat’ in the laboratory, has gained more attention recently because the technology may contribute to sustainable food production system. Currently, we face on food crisis with population growth, such as rapidly increasing demand for livestock products. Raising livestock becomes an environmental stressor since it requires large amount of natural resources and brings about 14.5% of total anthropogenic greenhouse gas emissions (Charles et al., 2019; Gerber et al., 2013; Grossi et al., 2019). Therefore, establishment of sustainable food resource production system for next generation is effective for environmental and ecological protection (Lynch & Pierrehumbert, 2019; Tuomisto, 2019). In addition, it is expected that food wastes are reduced and recycled in an effective manner (Lemaire & Limbourg, 2019).

Fish has been consumed increasingly worldwide, resulting in the enormous impacts on the ecosystem such as overfishing. In the present study, we focus on the developmental technology which creates ‘fish clean meat’ from fish cells for the first time in the world, in order to overcome the challenges in Sustainable Development Goal (SDG) 14: Conserve and sustainably use the oceans, seas and marine resources for sustainable development (FAO, 2015). We especially focused on fish fins and scales because 1) they are often discarded in the meal and 2) collecting the scales and fins partially does not take a life in fish. We believe that technology of creating ‘aquatic clean meat’ from fish brings potential benefits including environmental friendliness, animal welfare and sustainability.

Fish have a high level of ability in regeneration. They are able to regenerate various body parts such as fins which are equivalently a hand and foot in humans, hearts, neurons and so on. In general, regeneration occurs by dedifferentiation and/or self-renewal with basal stem cells (Jopling et al., 2010; Kikuchi et al., 2010; Sada et al., 2016; Sehring & Weidinger, 2020). Individual fish enables to regenerate their partially lost fins within a few weeks. What about the fins cut from the body, on the other hand? The cultured fin tissues cut from fish body shows explants around the tissues. However, the fin tissues have never been shaped to a complete fin as it was in culture flask. Then, what ability or characteristics have those cells which gathered for regeneration retained? If the cells have de-differentiated, what is the differentiation potential do they have?

In general, ES cells and the inner cell mass of early embryos as well as amphibian undifferentiated embryo cells are cultured without adding specific stimulating factors when they are differentiated and cultured in vitro, that is, serum, growth factors, etc. It is known to lead neural differentiation without external signals (Kamiya et al., 2011). Thus, basal state of undifferentiated cells had a fate to neural induction. Shimada *et al.* (2013) reported that the fins were derived from somatic mesoderm which has developed from somatic stem cells.

These results suggests the possibility that the fin cells in regeneration process was dedifferentiated and become undifferentiated cells, they may become basal neural somatic stem cells. If differentiated fish cells have the ability to differentiate into various types of cells such as muscle, fat, and fibers and nerves, they will be able to transform into cultured ‘aquatic’ clean meat.

## Materials and method

### Animal

Live thread-sail filefish *Stephanolepis cirrhifer* was captured near Jogashima Island in Yokosuka City, Kanagawa Prefecture, Japan. Specimens were maintained in aerated artificial seawater and its salinity and water temperature were maintained at 34.0 ppt and 20±2°C, respectively.

### Preparation of the cell line

The cell line of the thread-sail filefish *S. cirrhifer* was prepared the cells as follows. We obtained a 5 mm^2^ cut of fin tissue from the thread-sail filefish and placed it in 70% ethanol. The tissue was then washed three times with phosphate buffered saline (PBS), then six times with penicillin and streptomycin (MP Biomedicals, Santa Ana, CA) on ice. The dorsal fin tissue was cut into 1 mm squares in 0.25% Trypsin (MP Biomedicals)-0.02% EDTA (MP Biomedicals) solution and incubated at room temperature for 20 min. The tissue was centrifuged at 1100 rpm and washed twice with Leibovitz’s L-15 culture medium (Life Technologies, Carlsbad, CA). The tissue was immersed in L-15 media containing 10% fetal bovine serum (FBS) (Biowest, Nuaillé, France) and 1% Zell Shield (Minerva Biolabs, Berlin, Germany), then seeded in a 25 cm^2^ Collagen I coated flask (Thermo Fisher Scientific, Waltham, MA), and cultured in an incubator at 25°C. After 24 h, we confirmed the outgrowth surrounding the tissue. A couple of days later, the cells around the tissue were removed with TrypLE Express (Life Technologies), and seeded into a new flask. The culture media was replaced every 3 d and the cells were subcultured at 4.0×10^5^ cells/ml five times to establish the cell line, named. TSF Cell morphology was observed using an inverted microscope CKX41 (Olympus, Tokyo, Japan) connected to a digital camera ARTCAM-300MI-WOM (Artray, Tokyo, Japan).

### Induction of Differentiation

Fin cells from established *S. cirrhifer* cell line were evaluated their differentiation under various culture conditions. 4.0×10^5^ cells were seeded in L-15 media containing 10% FBS in a 25 cm^2^ Collagen I coated flask or Non-coated flask (Thermo Fisher Scientific). 24 h later, the cells were washed with PBS and treated with different media. For those seeded in Collagen I coated flask, AIM V Medium (Thermo Fisher Scientific) + 10% FBS, AIM V Medium (Thermo Fisher Scientific) only, L-15 + 10% heat-inactivated SeaGrow (EastCoast Biologics, North Berwick, ME), KBM Neural Stem Cell medium (Kohjin Bio Co. Ltd., Saitama, Japan) + 1X Neural Induction Supplement (Thermo Fisher Scientific). For the cells seeded in Non-coated flask, L-15 only. Cell differentiation was photographed every 60 seconds and the time-lapse movies were created with Axio Vision ver. 4.8 software (Carl Zeiss).

### Lipid staining

The fin cells were fixed in 4% paraformaldehyde (Wako Pure Chemical Industries, Ltd., Osaka, Japan) in phosphate-buffered saline (PBS) at RT for 1 h. The fixed cells were subjected to BODIPY® staining with Adipocyte Fluorescent Staining kit (Cosmo Bio Co. Ltd., Tokyo, Japan) following to manufacture’s protocol. For Oil Red O staining, the fixed cells were treated with 60% isopropanol for 5 min. After the isopropanol was removed, the cells were incubated with Oil Red O solution at RT for 25 min. The treated cells were observed with the microscope as stated above.

### Immunofluorescence

Differentiated cells were fixed in 4% paraformaldehyde (Wako Pure Chemical Industries, Ltd.) in PBS at RT for 12 h. The cells were incubated with Milli-Mark™ FluoroPan Neuronal Marker (Mouse IgG conjugated with Alexa 488) (Merck Millipore, Burlington, MA) as the manufacturer's instruction. The fluorescent cells were observed with a Zeiss Axio Observer. D1 microscope (Carl Zeiss, Oberkochen, Germany). An AxioCam HRc camera (Carl Zeiss) was used to photograph the images, and the images were analysed using AxioVision ver. 4.8 software (Carl Zeiss).

### Scanning electron microscopy (SEM)

The adherent cells on the collagen I coated slide were fixed with 0.4 ml of 2.5 % glutaraldehyde in culture medium in a 2 ml culture chamber overnight at 4 °C. The cells were postfixed with 2.0 % osmium tetroxide dissolved in PBS for 2 hours at 4 °C, and then were dehydrated in a graded series of ethanol, and freeze dried in a freeze drier VFD-21S (Vacuum device Inc., Ibaraki, Japan). They were coated with osmium using a POC-3 osmium plasma coater (Meiwafosis Co. Ltd., Tokyo, Japan) and observed in a field emission scanning electron microscope JSM-6700F (JEOL, Ltd., Tokyo, Japan) operated at 5 kV.

### Gas chromatography

Fatty acids in the fin, the fin cells and the differentiated cells were determined by gas chromatography. The extract sample (1 μl) was injected into a gas chromatograph 7890A (Agilent Technologies Inc., Santa Clara, CA) equipped with an Omega wax 320 30 m ×0.32 mm.

## Results

### Does the regeneration of fish fin tissue occur?

YES, the fin of *S. cirrhifer* began to regenerate soon after being cut. Basic structure of the fin ray was formed within two weeks and recovered to an original state four weeks later (Fig.1). On the other hand, a piece of the fin tissue which was cut apart from the fish body showed populated swarmer cells in the culture media (Fig. 2). The cell was about 20-50 μm in size. Subculturing was performed subsequently, and stable fibroblast-like cells were obtained at 5^th^ passage (Fig. 3a, Movie 1). We named the fibroblast-like cells as deSc (<u>de</u>differentiated *<u>S</u>tephanolepis <u>c</u>irrhifer*). To verify a chromosomal mutation had not occurred in the deSc cells, Q-banding stain analysis was performed and compared to wildtype *S. cirrhifer* which had been reported by Murofushi *et al*. (1980). As a result, the number of chromosomes was 33 (2n=30+X_1_X_2_Y) in 96% of deSc cells, which was identical to the wildtype *S. cirrhifer* (Supplementary Fig. 1). 4% of deSc cells showed 66 chromosomes since they might be in the middle cell division with replicated DNA.

**Figure 1.**
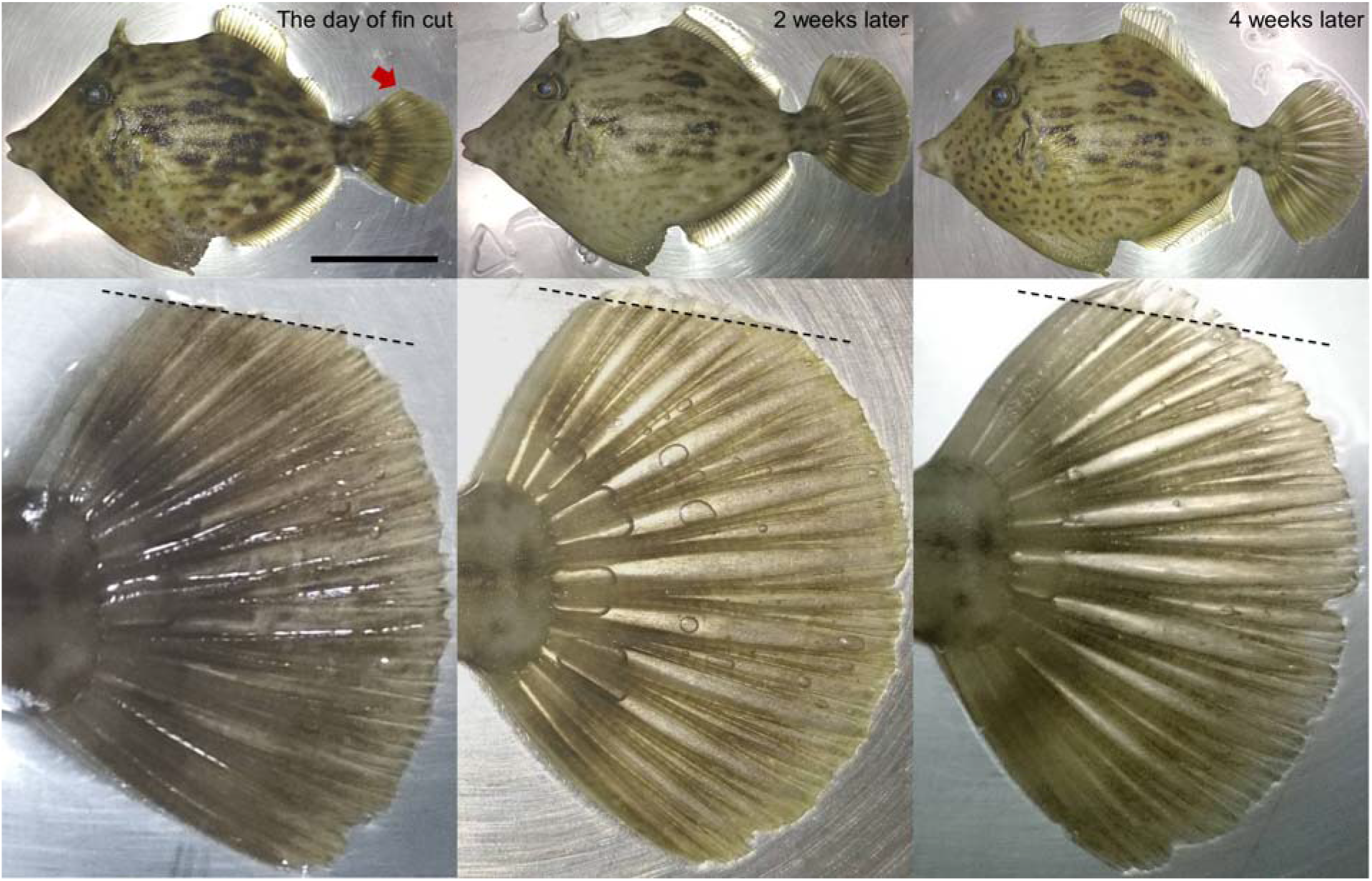
Regeneration of the fin in thread-sail filefish *Stephanolepis cirrhifer*. Upper; whole fish, lower; enlarged view of the red arrow part. Scale bar; 5 cm.

**Figure 2.**
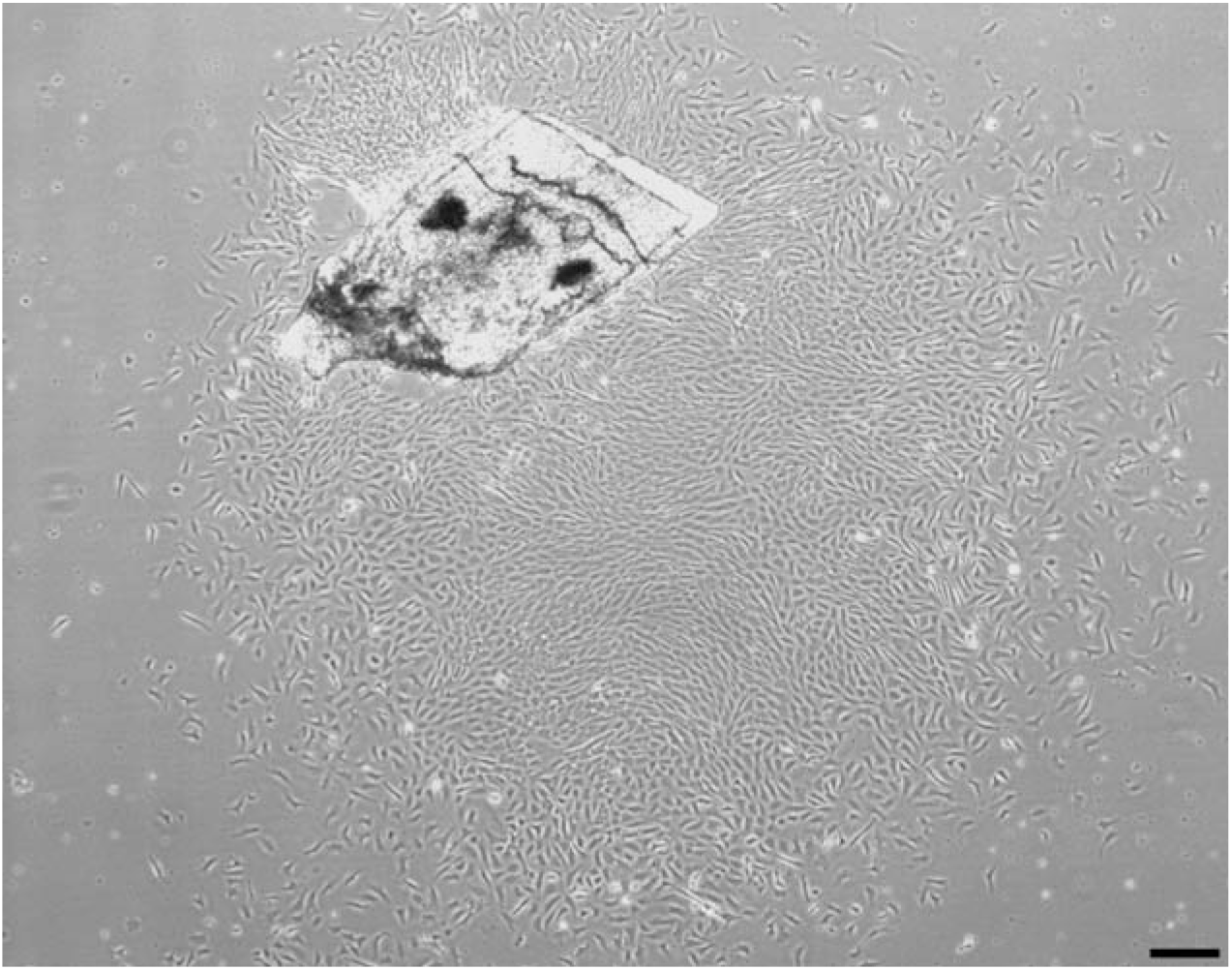
Swarmer calls from fin tissue of *S. cirrhifer*. Scale bar; 100 μm.

**Figure 3.**
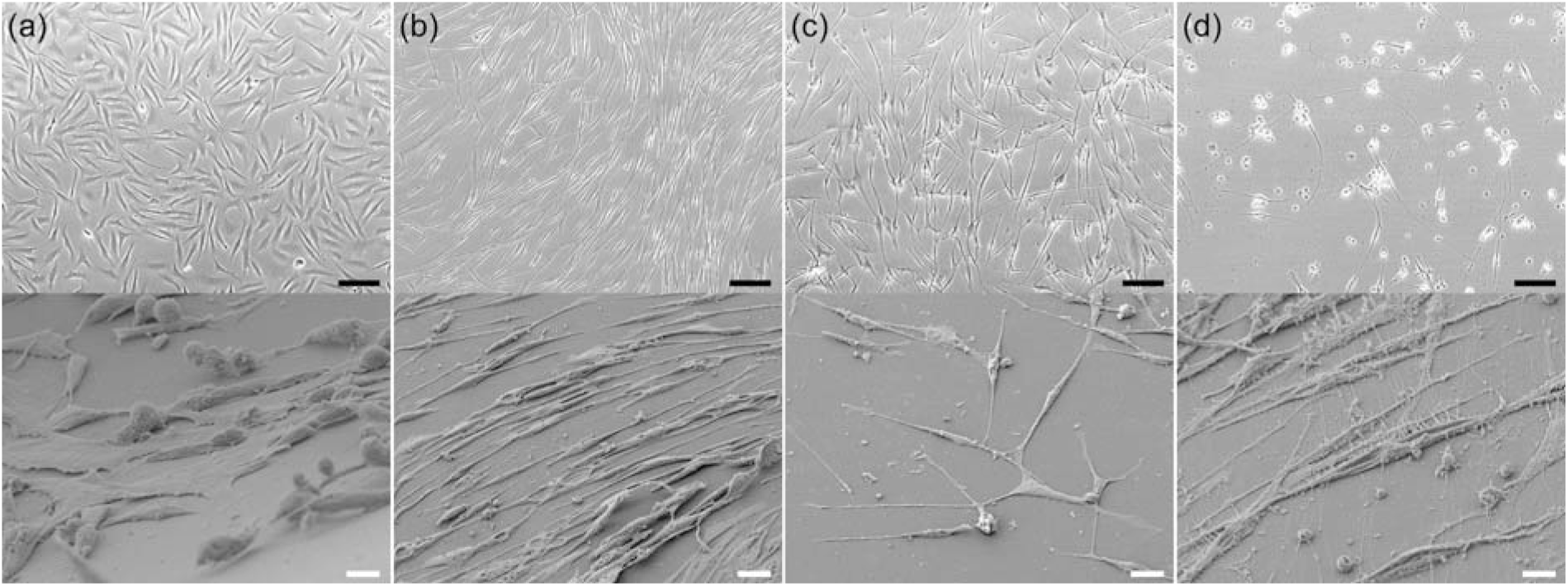
deSc cells differentiated various cell types. a) Normal fibroblast-like cells, b) Skeletal muscle-like cells, c) neural-like cells, d) neurofilaments. Upper; bright field image, scale bar; 50 μm. Lower; SEM image, magnification; ×1000 magnified, scale bar; 10 μm.

### Do the fin cells differentiate?

YES, they do. They differentiate to many types of cells indeed! To explore an optimal culture condition for the deSc cells, we had examined several culture media, serum, and extracellular matrix (ECM) and discovered very interesting characteristics (Table 1). The cell changed their morphology variously based on the combinations of those culture media, serum and ECM (Table 1, Fig. 3b, Movie 2). Within the various results, we focused on the neural-like cells which differentiated own morphology under culturing without serum (L-15 medium only) in a non-coated flask (Fig. 3c, Movie 3).

**Table 1.** Differentiation of deSc cells under various combinations of culture medium, serum, flask and additive.

| Cell Morphology | Culture Medium | Serum | Flask | Additive |
| --- | --- | --- | --- | --- |
| Fibloblast-like | L-15 | FBS | Collagen I-coated |  |
| Skeletal muscle-like | AIM V | FBS | Collagen I-coated |  |
| Neuronal-like | L-15 | — | Non-coated |  |
| Neurofilament | KBM | FBS | Collagen I-coated | Neural induction supplement |
| Adipocyte | L-15 | SeaGrow | Collagen I-coated |  |
| Spheroid | L-15 | FBS | Spheroid |  |
| Cocoon | L-15 | Horse | Non-coated |  |

### Is there differentiation potency in deSc cells?

YES, there is. Basal state of ES cells in the absence of neural differentiation inhibitors such as serum and transcription factor adopted neural fate (Kamiya et al., 2011). The present deSc cells which were cultured in the same condition have also differentiated to neural-like cells within 24h. The result led us to speculate that the deSc cells have potential of neural differentiate likewise if the cells had acquired pluripotency through dedifferentiation process. To demonstrate the hypothesis, we first attempted to induce neural differentiation directly with KBM Neural Stem Cell medium (Kohjin Bio Co., Ltd.) and Neural Induction Supplement (Thermo Fisher Scientific). As the result, neurofilaments were formed with 465 μm maximum length, and 45.71 μm/h average elongation speed (Fig. 3d, Movie 4, Table 1). Neural immunofluorescence suggested that those fin cells were virtually differentiated to the neural cells (Fig. 4). These results demonstrated that the deSc cells possess the characteristic of practicable direct-differentiation only with culture medium components. We have succeeded to induce the neural differentiation, which is a basal state of stem cells, by both the presence/absence of serum and the direct differentiation. We then came up with a question, “What will happen to the differentiation if serum is different?”

**Figure 4.**
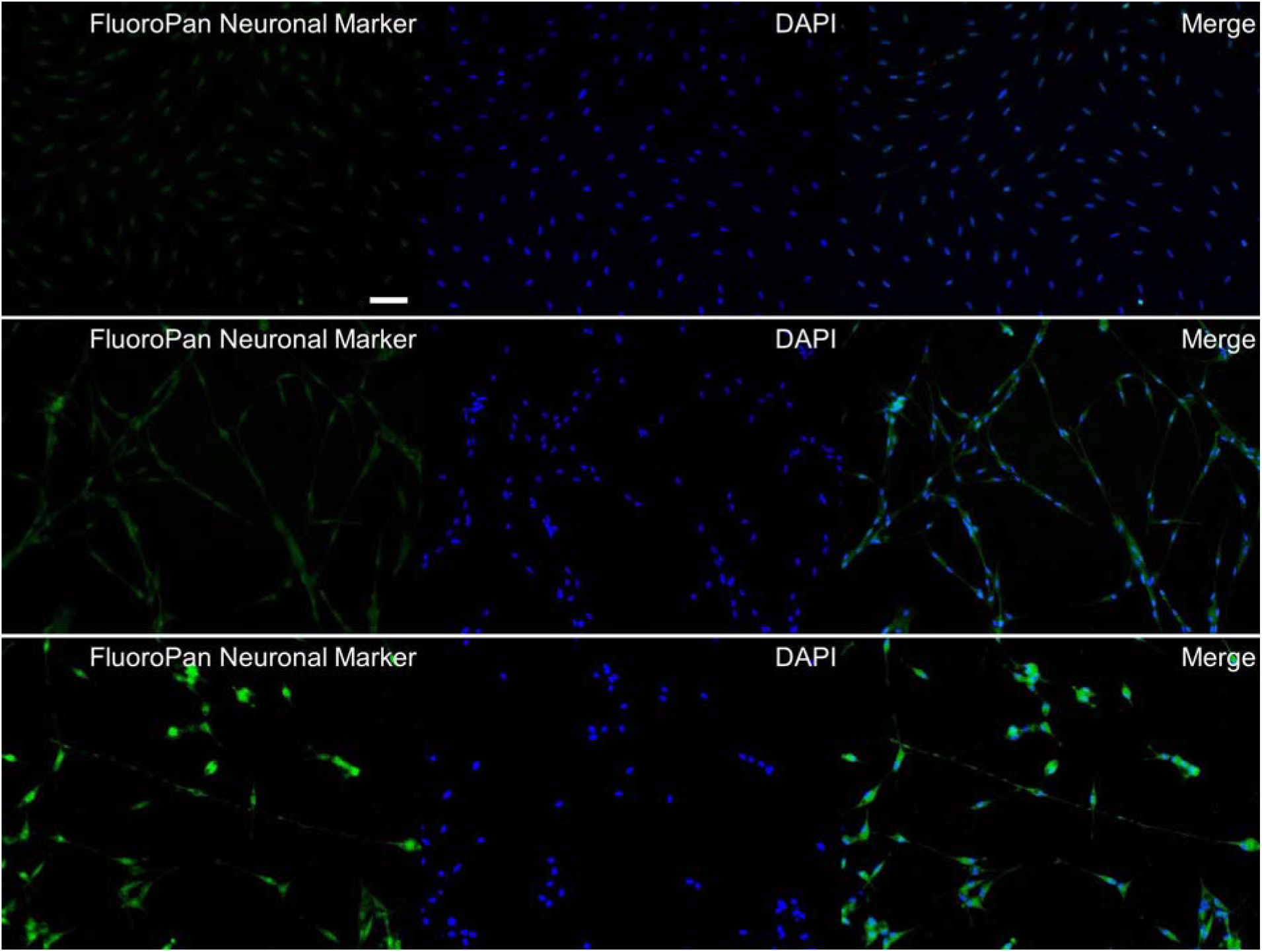
Immunofluorescence of neural deSc cells. Upper; normal deSc cells, middle; neural-like deSc cells, lower; neurofilaments. Scale bar; 50 μm.

### What happens to the cell differentiation when the serum is changed?

Very interesting phenomena were observed. In human iPS cells, culturing with different mammalian sera is reported to affect cell proliferations, differentiations, gene expressions and a stability of transcriptomes (Gstraunthaler, 2003; Shahdadfar et al., 2005). We next examined cell differential morphology under culturing with different sera. First, we assessed salmon serum SeaGrow which was the same taxonomical group as the deSc cells. Granule-like particles were appeared intracellularly five hours after the stimulation with the fish serum. Another three hours later, the cell morphology changed to the round, larger in size, which were adipocyte-like cells with 0.5-2.0 μm white droplets (Fig. 5a, Movie 5). In order to examine the white droplets for details, the cells were stained with Oil Red O and BODYPY, respectively, and also analyzed with a gas chromatography. The results showed that those white droplets were fat droplets (Supplementary Fig. 2). Therefore, the replacement to salmon serum in the culture media resulted in the differentiation to adipocyte in deSc cells.

**Figure 5.**
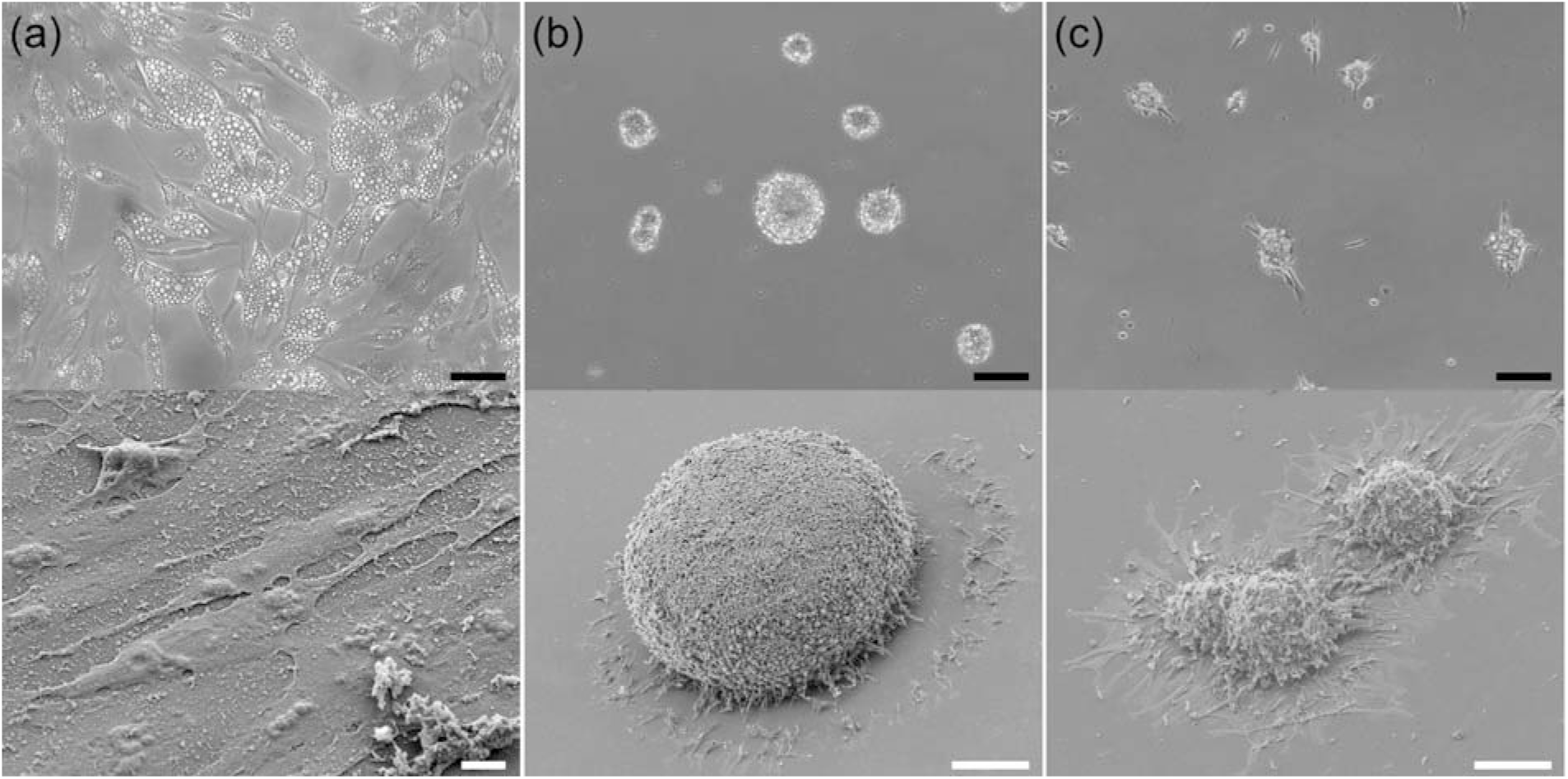
Differentiation of deSc cells to adipocyte (a), spheloid (b), and cocoon (c). Upper; bright field image, lower; SEM image. Scale bars; 50 μm (a) upper, 10 μm (a) lower, 100 μm (b) (c).

We next examined several mammalian sera other than FBS in the cell culture. With a rabbit and a sheep serum, the deSc cells did not survive, or did not present stable and uniform differentiation states (data not shown). However, we have witnessed an extraordinarily interesting phenomenon when examining a horse serum.

We, organisms consist of various tissues such as bones, cartilages, muscles, skin, which are established on living-scaffolds. In this point, the process of turning differentiated cells into a tissue body is significantly important. Recently, a three-dimensional spheroid has been reported as an important *in vitro* model in terms of similar functions and organization to biological tissues (Achilli et al., 2012). The deSc cells formed a spheroid with three-dimensional culture, which mammalian cells did likewise. (Fig. 5b, Movie 6). The size of spheroids were between 20 and 300 μm. Spheloids were formed by intaking their surrounding cells. Interestingly, stimulating the cells with the horse serum showed colonies which looked like suspended cell aggregates with adherent “feet” (Fig. 5c, Movie 7). We had named those colonies, “Cocoon.” The cocoons “walked” around in the culture flask at 38.46 μm/h average speed and also they moved largely every 3.5 h with absorbing or fusing surrounding other cocoons (Movie 8). Furthermore, single deSc cells which had not formed the cocoon or had separated from it showed the same behavior as the cocoons. Interestingly, ‘the independent cells’ which had not constituted the colonies showed neither a cell division nor cell proliferation. The colonies showed various sizes between 20 and 1000 μm diameter and the largest ones were able to be visualized macroscopically. Moreover, our observations confirmed that the colonies were stable in culture at least three weeks, as seen in spheroids culture too (data not shown).

To date, we are not able to specify what characteristics the cocoons acquired and what they differentiated to. However, they showed surprisingly dissimilar behaviors by acquiring the 3D structure with the legs compared to the normal cells. The cell differentiations mentioned above; skeletal muscle-like, adipocyte, spheloid and cocoons, were reversible processes by turning back to the basic culture condition; L-15 medium with 10% FBS in collagen-I coated flask, except the neural differentiation. Dedifferentiation which observed in the regenerating fin at an individual level was continuously observed at cellular levels too in fish. Moreover, the differentiation processes were surprisingly simple since its triggers were the culture media, sera and ECM.

## Discussion

Dedifferentiation mechanism is relatively a common phenomenon observed as a regeneration process in organisms. Our present study reports for the first time in a cellular level that the fish cells sampled during the regeneration process have a stem cell-like characteristic. We demonstrated that deSc cell differentiations were induced by “simple stimulus” such as medium, serum and ECM, without a specialized technique like a gene transfer. While the present studies focused on the thread-sail filefish *S. cirrhifer*, the differentiate phenomena in two-dimension have been observed in other several fish cells, too, more or less with differentiation potential (Supprementary Fig. 3). To date, sophisficated biological technology, gene transfer, enables to create tissues from various kinds of cells including iPS and ES cells (Karagiannis et al., 2019). Moreover, some research groups succeeded to produce other cells directly from fibroblast without going through those stem cells (Wada et al., 2013). Our studies suggested that the potential for ‘simple and direct differentiation’ is also commonly present in fish as well as in mammals.

We believe that finding such potential in edible fish is extremely significant because it is interesting to utilize the fins which are usually discarded as food wastes, effectively to the next-generation food processing technologies, including clean meat. Especially in fish meals, taste and texture of the preferment parts in favorite kinds of fishes will be able to be reproduced anytime without being affected by the seasonal catches. Moreover, aquatic clean meat has a merit of unnecessity of microplastic pollutions.

## Postscript

We would like to discuss the potential of fish cell research for its future development. Swann et al. (2014) reported that mouse-derived thymic epithelial cells generated a fish-like thymus under coexpression of *Foxn4* and *Foxn1*. Kherdjemil et al. (2016) and Nakamura et al. (2016) found that fish fin rays and fingers in mammals emerge from the same group of embryonic cells. Fish is known to have high homology to humans (Aparicio et al., 2002; Howe et al., 2013). These studies may suggest as follows; it is not impossible that cells can be utilized universally beyond species by resetting cell functions with the reprogramming technology; in other words, various types of mammalian cells can be created from fish cells. Moreover, our study shows the cocoon, the 3-D structured cells, which was interesting phenomenon. The cocoons use their legs like tentacles, which look like they have acquired a will, however, it is difficult to explain what the behavior and the legs are for at present. Re-creating organisms from cultured cells is hard to imagine, but interesting.

Organisms have acquired a variety of lifestyle from ocean to land by changing their fins to legs through evolution (Flammang et al., 2016). Fish fin is adopted and induced phenotypic changes in response to environmental changes (Standen et al., 2014). We demonstrated that deSc cells, the fins in cellular level, showed facile differentiation. These results and reports lead us an important factor – “serum,” that is cells are possible to make themselves change with the blood constituents. This is enormously interesting in terms of discussing evolutionary aspect in biodiversity because differentiation is significant phenomenon in evolution. At present, a mainstream approach to induce differentiation in scientific research is a DNA modification by gene transfer. However, it is hard to imagine those modifications had occurred naturally in ancient times. Then, what was involved with the differentiation other than the environmental gene alteration? What if a blood was an inducer of the differentiation? Since organisms often contact with blood in nature through predation etc., can’t we negate the possibility that the blood has been involve and served in biodiversity?

Finally, we came up with another question through our studies. Just as there are individual differences among living things, cells also have characteristics individually. However, every cell differentiates almost identically upon differentiation process. Could there be some cells that are difficult to differentiate? This is very mysterious point. In recent years, it has been reported that pluripotent stem cells are inherently present in living organisms in mesenchymal cells such as fibroblasts and bone marrow stromal cells in mammals (Kuroda et al., 2010). What if there is a leader cell which plays a role ordering a command to the other cells even among cells that look the same? When the leader cell is stimulated somehow, is a command issued so that the rest of all cells in the flask begin to change? To support our hypothesis, a behavioral coupling mechanism between neighboring epidermal stem cells was reported, that is, differentiation of a single stem cell was followed by the division of local neighboring mouse cells (Mesa et al., 2018). Examining biological cells of not only mammals but also various species will bring us greater knowledge of cell biological function in the future.

## Funding

This research received no specific grant from any funding agency in public, commercial or not-for-profit sectors.

## Acknowledgement

We would like to thank Mr. Tomohisa Ogawa for technical support.

## Supplemental Movie

Movie 1. Proliferation of normal deSc cells.

Movie 2. Normal deSc cells differentiate to skeletal muscle-like cells.

Movie 3. Normal deSc cells differentiate to neural-like cells.

Movie 4. Normal deSc cells differentiate to neurofilaments.

Movie 5. Normal deSc cells differentiate to adipocyte.

Movie 6. Normal deSc cells differentiate to spheroids.

Movie 7. Normal deSc cells differentiate to cocoons.

Movie 8. Normal deSc cells differentiate to cocoons.

## Supplementary Figure

Supplementary Figure 1. Chromosome of deSc cells, *Stephanolepis cirrhifer*, 2n=30+X_1_X_2_Y(33). Scale bar; 5 μm.

Supplementary Figure 2. Adipocyte staining with a) BODIPY and b) Oil-red O. Scale bar; 50 μm. c) gas chromatogram of normal deSc cells (upper) and adipocyte (lower).

Supplementary Figure 3. Differentiation of scorpion fish, *Sebastiscus marmoratu*s. a) normal, b) neural-like cells under bright-field. Scale bar; 50 μm.

## Notes

#### Summary of Updates

Some errors in References section are revised.

## References

Achilli TM, Meyer J, Morgan JR. (2012). Advances in the formation, use and understanding of multi-cellular spheroids. Expert Opin Biol Ther. 12:1347–60.

Aparicio S, Chapman J, Stupka E, Putnam N, Chia JM, Dehal P, Christoffels A, Rash S, Hoon S, Smit A, Gelpke MD, Roach J, Oh T, Ho IY, Wong M, Detter C, Verhoef F, Predki P, Tay A, Lucas S, Richardson P, Smith SF, Clark MS, Edwards YJ, Doggett N, Zharkikh A, Tavtigian SV, Pruss D, Barnstead M, Evans C, Baden H, Powell J, Glusman G, Rowen L, Hood L, Tan YH, Elgar G, Hawkins T, Venkatesh B, Rokhsar D, Brenner S. (2002). Whole-genome shotgun assembly and analysis of the genome of Fugu rubripes. Science 297:1301–10.

Charles A, Kalikoski D, Macnaughton A. (2019). Addressing the climate change and poverty nexus: a coordinated approach in the context of the 2030 agenda and the Paris agreement. Rome. FAO

Food and Agriculture Organization of the United Nations (FAO). FAO and the 17 Sustainable Development Goals. United Nations, Rome (2015)

Flammang BE, Suvarnaraksha A, Markiewicz J, Soares D. (2016). Tetrapod-like pelvic girdle in a walking cavefish. Scientific Reports 6:23711.

Gerber PJ, Steinfeld H, Henderson B, Mottet A, Opio C, Dijkman J, Falcucci A, Tempio G. (2013). Tackling climate change through livestock: a global assessment of emissions and mitigation opportunities. Rome: FAO. Available from http://www.fao.org/3/a-i3437e.pdf

Grossi G, Goglio P, Vitali A, Williams AG. (2019). Livestock and climate change: impact of livestock on climate and mitigation strategies. Animal Frontiers 9:69–76.

Gstraunthaler G. (2003). Alternatives to the Use of Fetal Bovine Serum: Serum-free Cell Culture. ALTEX. 20:275–81.

Howe K, Clark MD, Torroja CF, Torrance J, Berthelot C, Muffato M, Collins JE, Humphray S, McLaren K, Matthews L, McLaren S, Sealy I, Caccamo M, Churcher C, Scott C, Barrett JC, Koch R, Rauch GJ, White S, Chow W, Kilian B, Quintais LT, Guerra-Assunção JA, Zhou Y, Gu Y, Yen J, Vogel JH, Eyre T, Redmond S, Banerjee R, Chi J, Fu B, Langley E, Maguire SF, Laird GK, Lloyd D, Kenyon E, Donaldson S, Sehra H, Almeida-King J, Loveland J, Trevanion S, Jones M, Quail M, Willey D, Hunt A, Burton J, Sims S, McLay K, Plumb B, Davis J, Clee C, Oliver K, Clark R, Riddle C, Elliot D, Threadgold G, Harden G, Ware D, Begum S, Mortimore B, Kerry G, Heath P, Phillimore B, Tracey A, Corby N, Dunn M, Johnson C, Wood J, Clark S, Pelan S, Griffiths G, Smith M, Glithero R, Howden P, Barker N, Lloyd C, Stevens C, Harley J, Holt K, Panagiotidis G, Lovell J, Beasley H, Henderson C, Gordon D, Auger K, Wright D, Collins J, Raisen C, Dyer L, Leung K, Robertson L, Ambridge K, Leongamornlert D, McGuire S, Gilderthorp R, Griffiths C, Manthravadi D, Nichol S, Barker G, Whitehead S, Kay M, Brown J, Murnane C, Gray E, Humphries M, Sycamore N, Barker D, Saunders D, Wallis J, Babbage A, Hammond S, Mashreghi-Mohammadi M, Barr L, Martin S, Wray P, Ellington A, Matthews N, Ellwood M, Woodmansey R, Clark G, Cooper J, Tromans A, Grafham D, Skuce C, Pandian R, Andrews R, Harrison E, Kimberley A, Garnett J, Fosker N, Hall R, Garner P, Kelly D, Bird C, Palmer S, Gehring I, Berger A, Dooley CM, Ersan-Ürün Z, Eser C, Geiger H, Geisler M, Karotki L, Kirn A, Konantz J, Konantz M, Oberländer M, Rudolph-Geiger S, Teucke M, Lanz C, Raddatz G, Osoegawa K, Zhu B, Rapp A, Widaa S, Langford C, Yang F, Schuster SC, Carter NP, Harrow J, Ning Z, Herrero J, Searle SM, Enright A, Geisler R, Plasterk RH, Lee C, Westerfield M, de Jong PJ, Zon LI, Postlethwait JH, Nüsslein-Volhard C, Hubbard TJ, Roest Crollius H, Rogers J, Stemple DL. (2013). The zebrafish reference genome sequence and its relationship to the human genome. Nature 496:498–503.

Jopling C, Sleep E, Raya M, Martí M, Raya A, Izpisúa Belmonte JC. (2010). Zebrafish heart regeneration occurs by cardiomyocyte dedifferentiation and proliferation. Nature 464:606–9.

Kamiya D, Banno S, Sasai N, Ohgushi M, Inomata H, Watanabe K, Kawada M, Yakura R, Kiyonari H, Nakao K, Jakt LM, Nishikawa S, Sasai Y. (2011). Intrinsic transition of embryonic stem-cell differentiation into neural progenitors. Nature 470:503–509.

Karagiannis P, Takahashi K, Saito M, Yoshida Y, Okita K, Watanabe A, Inoue H, Yamashita JK, Todani M, Nakagawa M, Osawa M, Yashiro Y, Yamanaka S, Osafune K. (2019). Induced pluripotent stem cells and their use in human models of disease and development. Physioligical Reviews 99:79–114.

Kherdjemil Y, Lalonde RL, Sheth R, Dumouchel A, de Martino G, Pineault KM, Wellik DM, Stadler HS, Akimenko MA, Kmita M. (2016). Evolution of Hoxa11 regulation in vertebrates is linked to the pentadactyl state. Nature 539:89–92.

Kikuchi K, Holdway JE, Werdich AA, Anderson RM, Fang Y, Egnaczyk GF, Evans T, Macrae CA, Stainier DY, Poss KD. (2010). Primary contribution to zebrafish heart regeneration by gata4(+) cardiomyocytes. Nature 464:601–5.

Kuroda Y, Kitada M, Wakao S, Nishikawa K, Tanimura Y, Makinoshima H, Goda M, Akashi H, Inutsuka A, Niwa A, Shigemoto T, Nabeshima Y, Nakahata T, Nabeshima Y, Fujiyoshi Y, Dezawa M. (2010). Unique multipotent cells in adult human mesenchymal cell populations. PNAS 107:8639–43.

Lemaire A, Limbourg S. (2019). How can food loss and waste management achieve sustainable development goals? Journal of Cleaner Production 234:1221–1234.

Lynch J, Pierrehumbert R. (2019). Climate impacts of cultured meat and beef cattle. Frontiers in Sustainable Food Systems 3:5.

Mesa KR, Kawaguchi K, Cockburn K, Gonzalez D, Boucher J, Xin T, Klein AM, Greco V. (2018). Homeostatic Epidermal Stem Cell Self-Renewal Is Driven by Local Differentiation. Cell Stem Cell 23:677–686.e4.

Murofushi M, Oikawa S, Nishikawa S, Yosida TH. (1980). Cytogenetical studies on fishes. III. Multiple sex chromosome mechanism in the filefish, *Stephanolepis cirrhifer*. The Japanese Journal of Genetics 55: 127–131.

Nakamura T, Gehrke AR, Lemberg J, Szymaszek J, Shubin NH. (2016). Digits and fin rays share common developmental histories. Nature 537:225–228.

Sada A, Jacob F, Leung E, Wang S, White BS, Shalloway D, Tumbar T. (2016). Defining the cellular lineage hierarchy in the interfollicular epidermis of adult skin. Nature Cell Biology 18:619–31.

Sehring IM, Weidinger G. (2020). Recent advancements in understanding fin regeneration in zebrafish. Wiley Interdiscip Rev Dev Biol. 9:e367.

Shahdadfar A, Frønsdal K, Haug T, Reinholt FP, Brinchmann JE. (2005). In vitro expansion of human mesenchymal stem cells: choice of serum is a determinant of cell proliferation, differentiation, gene expression, and transcriptome stability. Stem Cells 23:1357–66.

Shimada A, Kawanishi T, Kaneko T, Yoshihara H, Yano T, Inohaya K, Kinoshita M, Kamei Y, Tamura K, Takeda H. (2013). Trunk exoskeleton in teleosts is mesodermal in origin. Nature Communications 4:1639.

Standen EM, Du TY, Larsson HC. (2014). Developmental plasticity and the origin of tetrapods. Nature 513: 54–8.

Swann JB, Weyn A, Nagakubo D, Bleul CC, Toyoda A, Happe C, Netuschil N, Hess I, Haas-Assenbaum A, Taniguchi Y, Schorpp M, Boehm T. (2014). Conversion of the thymus into a bipotent lymphoid organ by replacement of FOXN1 with its paralog, FOXN4. Cell Reports 8:1184–97.

Tuomisto HL. (2019). The eco-friendly burger: Could cultured meat improve the environmental sustainability of meat products? EMBO Reports 20: e47395.

Wada R, Muraoka N, Inagawa K, Yamakawa H, Miyamoto K, Sadahiro T, Umei T, Kaneda R, Suzuki T, Kamiya K, Tohyama S, Yuasa S, Kokaji K, Aeba R, Yozu R, Yamagishi H, Kitamura T, Fukuda K, Ieda M. (2013). Induction of human cardiomyocyte-like cells from fibroblasts by defined factors. PNAS 110: 12667–72.

