## Supplementary figures and images for "Differentiation potential of fish fins –Effective utilization of the fins as food wastes–"

### Supplemental Figure 1

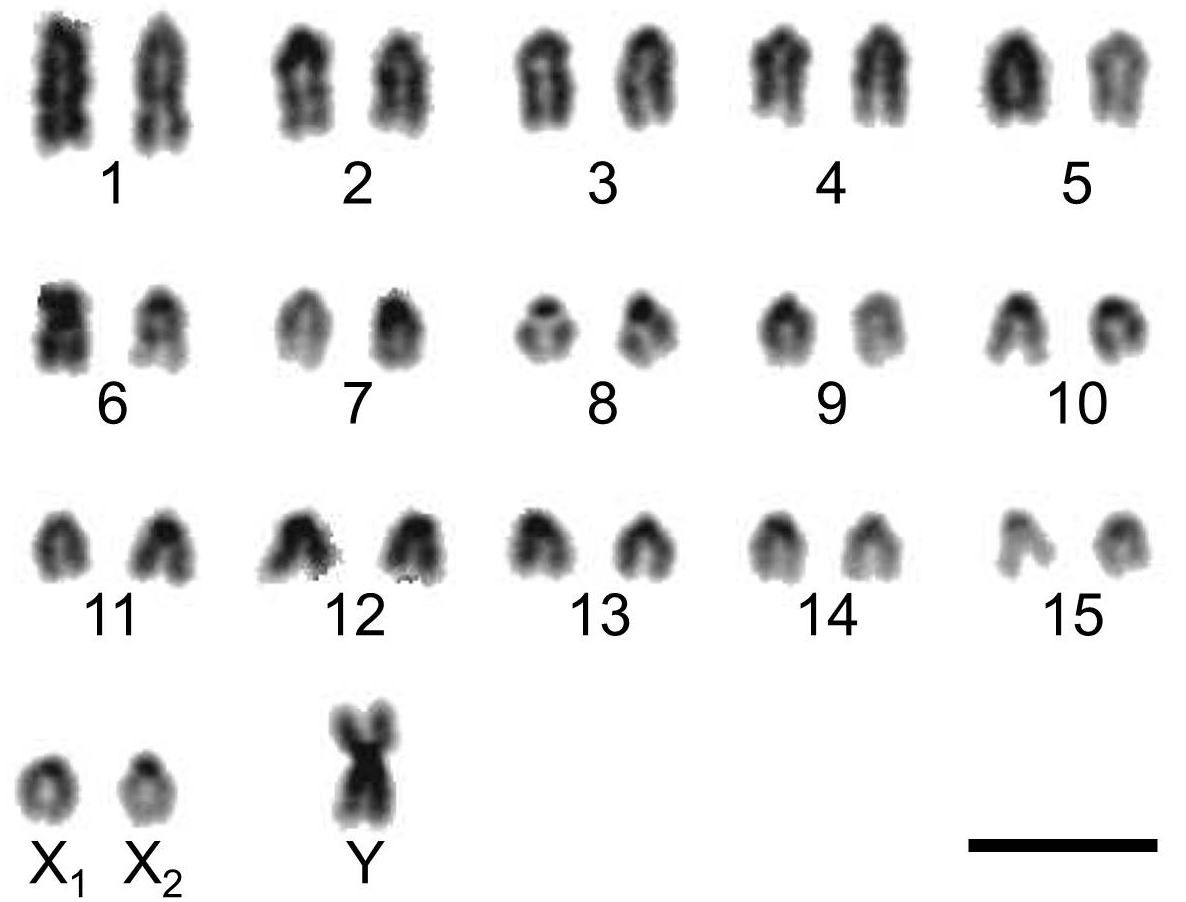

### Supplemental Figure 2ab

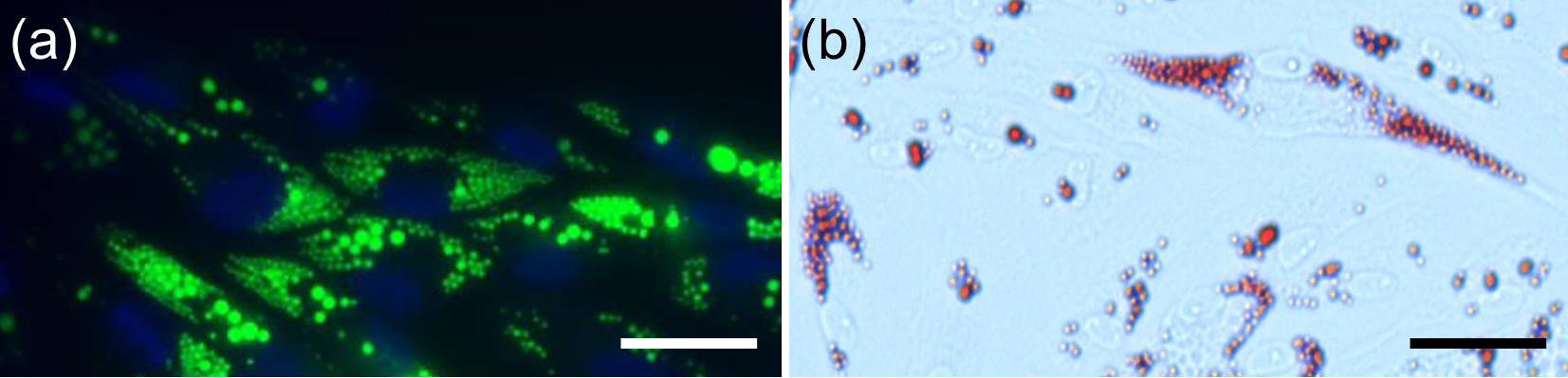

### Supplemental Figure 2c

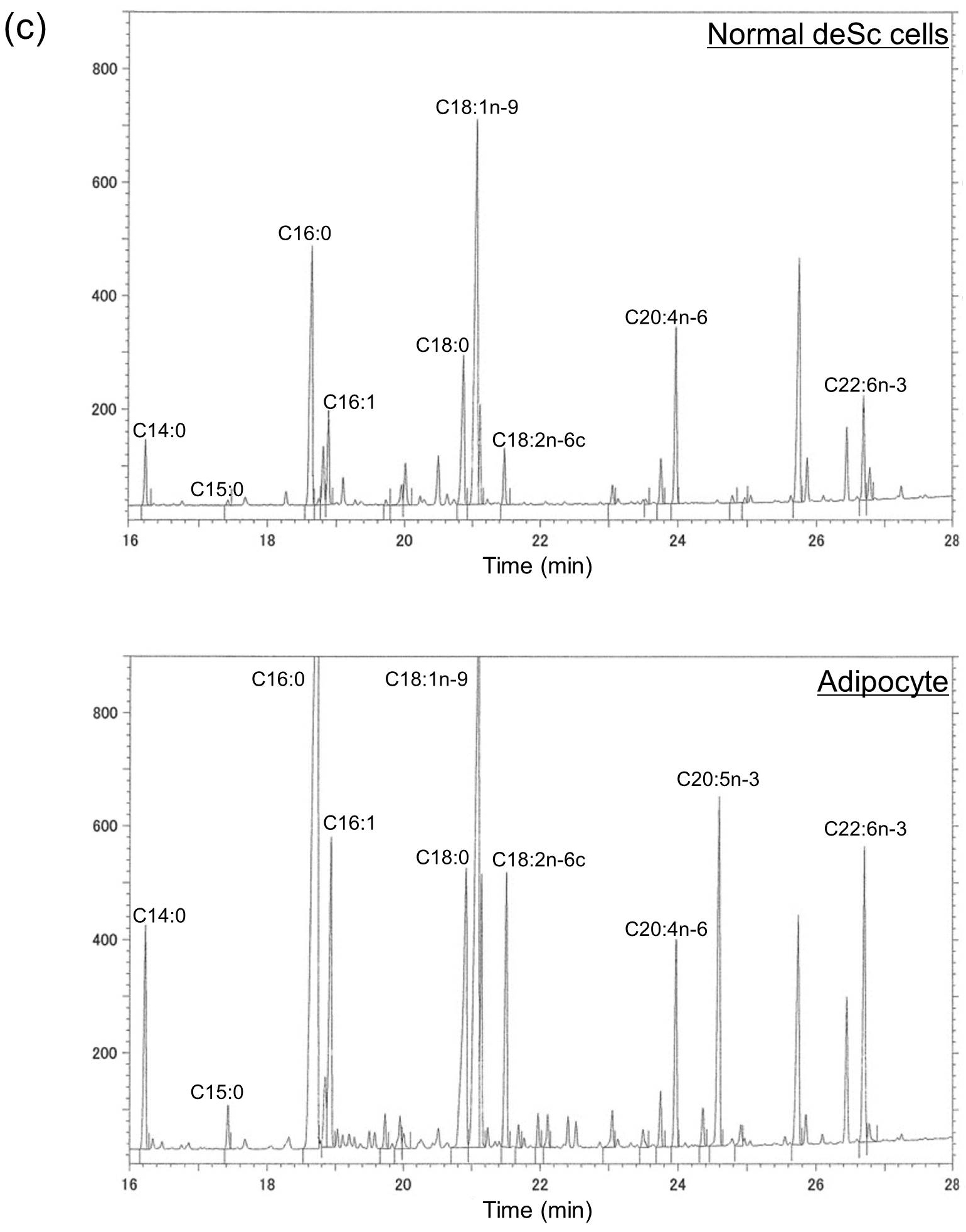

### Supplemental Figure 3

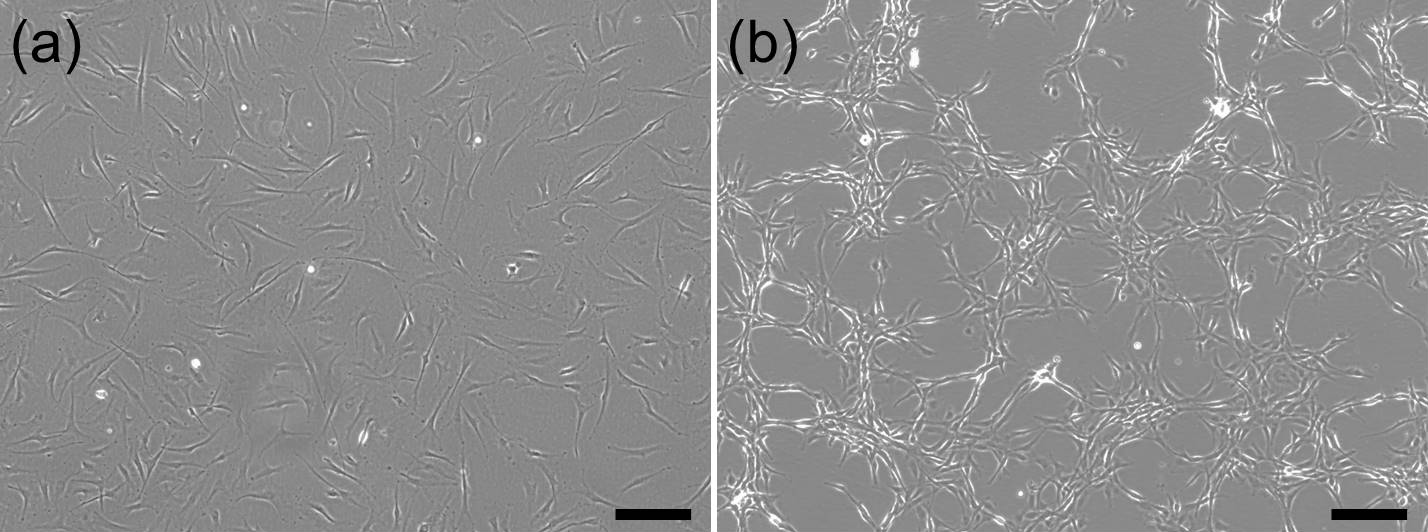
